# LongPolyASE: An end-to-end framework for allele-specific gene and isoform analysis in polyploids using long-read RNA-seq

**DOI:** 10.64898/2026.02.03.703480

**Authors:** Nadja Nolte, Kristina Gruden, Marko Petek

## Abstract

**Motivation:** Long-read RNA-seq and phased reference genomes enable haplotype-resolved gene and isoform expression analysis. While methods and tools exist for diploid organisms, analysis tools for polyploids are lacking.

**Results:** We developed an end-to-end framework for allele-specific gene and isoform analysis in polyploids with three components: Syntelogfinder identifies syntenic genes in phased assemblies; longrnaseq quantifies transcripts, discovers novel isoforms, and performs quality control of long-read RNA-seq; and PolyASE analyzes differential allelic expression, differential isoform usage between conditions, and structural differences in major isoforms between haplotypes. We demonstrate the use of the framework on diploid rice and autotetraploid potato.

**Availability and Implementation:** Syntelogfinder and longrnaseq are implemented in Nextflow and available on GitHub. PolyASE is a Python package available on PyPI. The framework is fully documented and tutorials are provided.

**Supplementary information:** Supplementary data are available online and on Zenodo.

## Introduction

Haplotype-resolved (phased) reference genomes reconstruct the distinct chromosomal copies (haplotypes) present in an organism separately rather than collapsing them into a single consensus. These genome assemblies are becoming available for a wide range of organisms, from personal diploid genomes of humans (Chin et al., 2016) to haplotype-resolved genomes of polyploid plant cultivars such as hexaploid wheat (Jia et al., 2023) and tetraploid potato (Sun et al., 2022). These phased reference genomes simplify expression analysis on haplotype level and reduce mapping bias in allele-specific expression (ASE) analysis (Degner et al., 2009).

Long-read RNA sequencing (RNA-seq) of full-length transcripts commercialized by Oxford Nanopore Technologies (ONT) and PacBio offers several key advantages over short-read RNA-seq: discovery of novel isoforms (Monzó et al., 2025), improvement of isoform quantification (Chen et al., 2025), and unique assignment of transcripts from highly similar loci to their origin based on single nucleotide variations. Combined with phased reference genomes, long-read RNA-seq enables straightforward quantification of allele-specific expression at both gene and isoform levels.

Allele-specific expression analysis can identify genes with imbalanced expression between haplotypes, revealing potential *cis*-regulatory differences, such as promoter variations or epigenetic modifications, that inhibit or increase gene expression from specific alleles. When comparing allelic expression across different conditions, e.g., developmental stages or environmental conditions, ASE analysis can identify *trans-*regulatory factors that differentially modulate haplotype expression in a condition-dependent manner, for instance condition-specific transcription factors (Metzger et al., 2016).

However, several challenges limit the application of phased assemblies and long-read RNA-seq for allele-specific gene and isoform expression analysis in polyploids. The majority of tools and pipelines for allele-specific expression analysis are developed for short-read RNA-seq and diploid organisms (Cleary & Seoighe, 2021; Mattevi et al., 2025). Current tools for long-read allele-specific gene and isoform analysis are phasing reads into maternal and paternal haplotype based on one haplotype reference genomes e.g. FLAIR2 (Tang et al., 2024), isoLASER (Quinones-Valdez et al., 2025), longcallR (Huang et al., 2025) or assigning reads based on alignments to phased genomes of diploid model organisms e.g. LORALS (Glinos et al., 2022). These tools lack options for analysis of more than two alleles and are therefore not applicable for polyploid organisms with phased assemblies. While few tools with polyploid support exist e.g. BYASE (Dong et al., 2020), they do not utilize phased references or long-read RNA-seq.

Phased reference genomes allow allele quantification without reference bias, but there are some limitations of their use. In phased reference assemblies, gene annotations are typically generated independently for each haplotype, resulting in non-unified gene identifiers and gene models across haplotypes. Therefore, syntelogs (Huynen & Bork, 1998), orthologous or paralogous genes that maintain their relative chromosomal position across haplotypes, must be identified before comparative expression analysis. Additionally, inconsistent gene annotation between haplotypes, commonly arising from length differences in the annotation of untranslated regions (UTR), can lead to biased haplotype level quantification (Nolte et al., 2025).

To address the gap in tools and the challenges of phased reference genomes for the analysis of haplotype-specific expression using long-read RNA-seq in polyploids, we developed an end-to-end framework for allele-specific gene and isoform expression analysis using long-read RNA-seq in diploid and polyploid organisms (Fig. 1). The framework consists of three components: (1) Syntelogfinder, a Nextflow (Di Tommaso et al., 2017; Langer et al., 2025) pipeline that groups genes by syntenic relationships and identifies length differences; (2) longrnaseq, a Nextflow pipeline for novel isoform discovery, haplotype level quantification, and quality control; and (3) PolyASE, a Python package for statistical testing and visualization of allelic imbalance, condition-dependent expression differences, differential isoform usage, and variation in major-expressed isoforms between haplotypes. We tested the framework on diploid rice and validated it on autotetraploid potato, using ONT long-read RNA-seq, identifying *cis*-regulatory variation, tissue-specific trans-regulatory effects, differential isoform expression of a novel identified gene linked to rice tillering regulation, and haplotype-specific splicing differences in potato.

**Figure 1:**
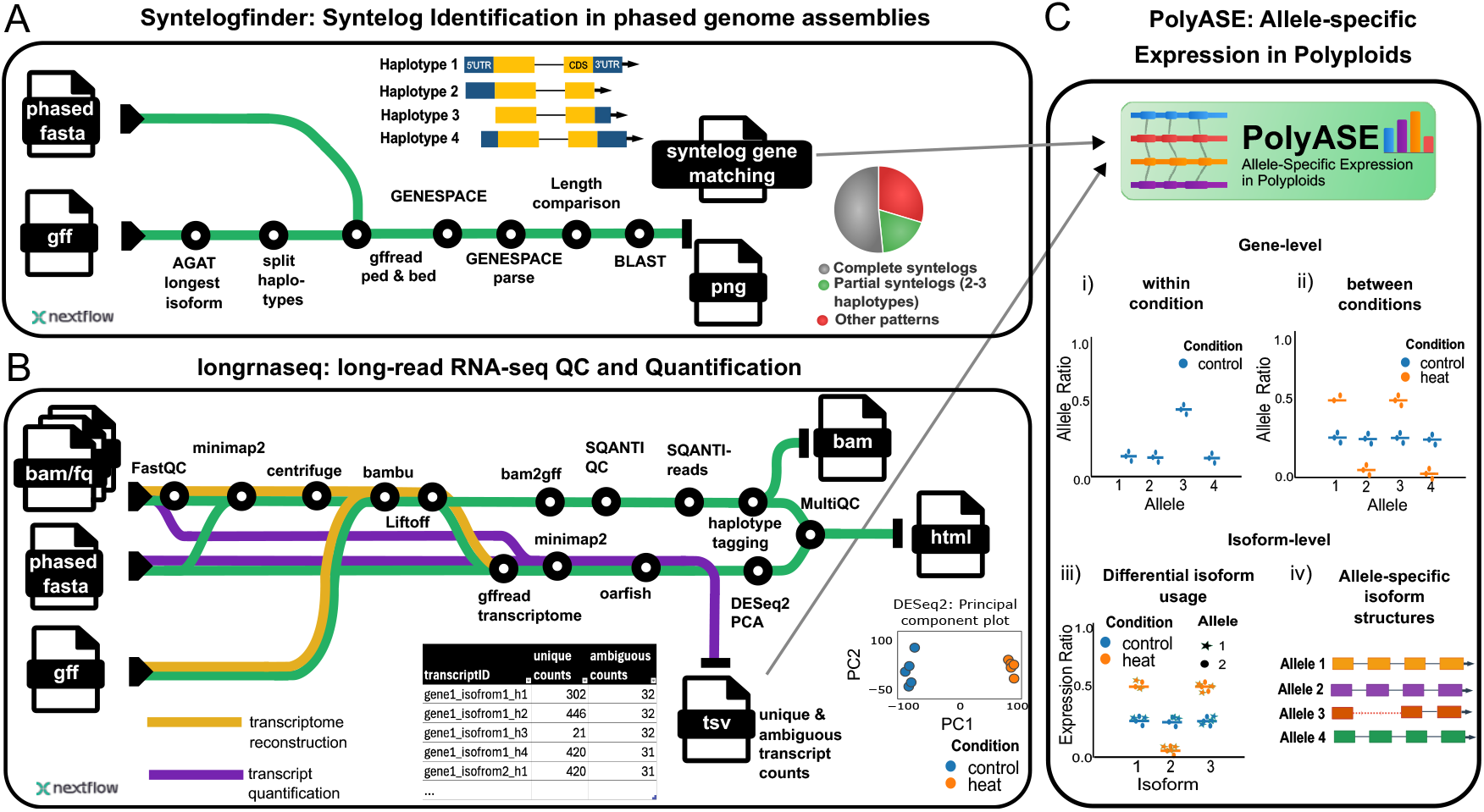
Overview of the allele-specific expression analysis framework for polyploid organisms using long-read RNA-seq. The framework consists of three integrated components: **(A) Syntelogfinder** identifies syntelogs across phased haplotypes using GENESPACE. Genes are categorized based on their copy number and syntenic relationships across haplotypes. Single-copy syntelogs, which are present on each haplotype with no gene duplications within any haplotype, are selected for allelic analysis. Steps in the pipeline are connected by a green line. **(B) longrnaseq** processes Oxford Nanopore Technologies (ONT) and PacBio long-read RNA data for discovery of novel isoforms and genes (yellow) and isoform level quantification (purple). First, the RNA-seq reads are aligned to a phased reference genome with minimap2, novel transcripts are discovered with Bambu, novel annotations are transferred across haplotypes with Liftoff, and transcripts are quantified with Oarfish. Further, quality control is performed with SQANTI3, SQANTI-reads, and Centrifuge, with comprehensive MultiQC reporting. Lastly, haplotype-tagged alignments enable visualization of allele-specific gene and isoform expression in genome browsers. **(C) PolyASE** performs statistical analysis and visualization of allele-specific expression patterns, including: (i) identification of genes with allelic imbalance within conditions (*cis*-regulatory variation), (ii) detection of condition-dependent changes in allelic ratios (*trans-*regulatory effects), (iii) differential isoform usage testing between conditions, and (iv) identification of structural differences in major isoforms between haplotypes.

## Methods and Implementation

### Pipeline Overview

We developed three integrated tools for analyzing allele-specific and isoform expression in polyploid genomes (Fig. 1): Syntelogfinder for identifying syntenic gene relationships, longrnaseq for processing long-read RNA-seq data, and PolyASE for downstream analysis and statistical testing.

### Identification of Syntenic Genes

The Syntelogfinder pipeline (Fig. 1 A) enables syntelog identification from phased reference genomes when gene annotations and a phased assembly are available. The pipeline splits annotations by haplotype, extracts the longest isoform per gene with agat_sp_keep_longest_isoform.pl v1.4.0 (Jacques Dainat et al., 2025), obtains protein sequences of protein-coding genes, and runs GENESPACE v1.3.1 (Lovell et al., 2022), which executes OrthoFinder2 (Emms & Kelly, 2019) and MCScanX (Wang et al., 2012). GENESPACE was configured with onlySameChrs=True to restrict syntelog identification to the same chromosome and ploidy=1 as genomes were separated into haplotypes, with one haplotype randomly designated as “refGenome”. The query_pangenes function identified gene groups with orthologous, paralogous, and syntenic relationships. Pangene output was parsed using a custom script to assign genes to groups based on haplotype presence and synteny. Single-copy syntelogs, i.e. those present on each haplotype with no gene duplications within any haplotype, were analyzed for sequence length differences and transcript similarity via BLASTn v2.16.0 (Altschul et al., 1990).

### Long-Read RNA-seq Processing and Quantification

The longrnaseq pipeline (Fig. 1 B) processes PacBio and ONT long-read RNA-seq data through two subworkflows: novel transcript discovery and quantification. Raw reads were quality-checked with FastQC v0.12.1 and aligned to phased reference genomes using minimap2 v2.29-r1283 (Li, 2018, 2021). ONT cDNA reads were aligned to the genome with parameters “-ax splice -N 200 --secondary=yes -G 10000 --secondary-seq”, and to the transcriptome with “--eqx -N 200 -ax map-ont”. PacBio reads were aligned with “-ax splice:hq -uf -N 200 --secondary=yes -G 10000 --secondary-seq” (genome) and “--eqx -N 200 -ax map-hifi” (transcriptome). For large genomes, the “-I 16g” option was added.

Novel transcripts and genes were identified from genome alignments using Bambu v3.0.8 (Chen et al., 2023) with default parameters. To mitigate haplotype-specific annotation bias, Bambu gene models were transferred between haplotypes using Liftoff (Shumate & Salzberg, 2021) with parameters “-infer_genes -a 0.7 -s 0.7 -copies -sc 0.7 -overlap 1.0”. Liftoff and original annotations were merged with gffcompare v0.12.7 (Pertea & Pertea, 2020) and identifiers were corrected with a custom script. Transcript quality control was performed with SQANTI3 v5.5.1 (Pardo-Palacios et al., 2024) with parameters “-min_ref_len 0 –skipORF” and SQANTI-reads v1.0 (Keil et al., 2024) with parameters “--force_id_ignore --pca_tables”.

Transcripts were extracted with gffread v0.12.7, aligned to transcripts with minimap2, and quantified using oarfish v0.9.0 (Zare Jousheghani et al., 2025) with parameters “--filter-group no-filters --write-assignment-probs --score-threshold 1.0”. Unaligned reads were screened for contamination using Centrifuge v1.0.4 (database build date: 03.03.2018). For visualization, aligned reads were subsampled using samtools view v1.21 (Danecek et al., 2021) and assigned a haplotype (‘HP’) tag based on the location of the primary alignment. This enables visualization of allele-specific expression and isoforms structures in genome browsers such as JBrowse2 (Diesh et al., 2023). Quality control reports were generated with MultiQC v1.25.1 (Ewels et al., 2016). The pipeline outputs transcript-level counts for uniquely and ambiguously mapping reads at the haplotype level, is modular and containerized following nf-core principles (Di Tommaso et al., 2017; Ewels et al., 2020; Langer et al., 2025), was tested on a SLURM-managed HPC cluster, and handles large genomes including phased wheat.

### Downstream Analysis of Allelic Expression

All downstream processing was implemented in PolyASE v1.3.0 (Fig. 1 C), a Python package for inspection of allelic counts and testing for imbalanced expression of syntenic genes, as well as isoform usage and structure. The necessary input files are transcript counts and grouping of genes based on syntenic relationships, which are generated by the two described Nextflow pipelines.

For gene-level analysis, transcript counts were aggregated by summing unique counts. For ambiguous counts, the mean across all transcripts per gene was used, assuming transcripts from the same gene multimap to the same genes. PolyASE visualizes allelic ratios and filters genes with high multimapping rates or length differences between haplotypes.

Multimapping rates were calculated for each syntenic group as: *gene ambiguity ratio* = (*total multimapping counts*) / (*total unique* + *multimapping counts*). This ratio is identical for all genes within a syntenic group and filters groups where allelic expression cannot be reliably tested for allelic imbalance due to similarity to other genes. Allelic ratios were calculated as: *alleic ratio per allele* = *unique counts per allele*/ *unique counts per gene*. For each transcript, the ambiguity ratio was computed as: *transcript ambiguity ratio* = (*multimapping counts per transcript*) / (*unique* + *multimapping counts per transcript*). Gene-level transcript ambiguity was calculated as a weighted average: 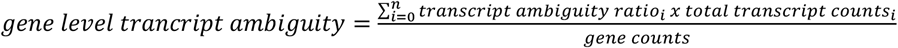. This metric filters genes unsuitable, due to high isoform similarity, for differential isoform usage testing.

### Statistical Testing

All tests used raw unique counts and a likelihood ratio test based on a beta-binomial model. For allelic imbalance within conditions, each allele was tested against the expected count ratio (1/ploidy × total gene counts per replicate). For comparing allelic ratios between conditions, unique counts from all replicates were compared. For differential isoform usage between conditions, unique counts per isoform were compared.

For multi-isoform genes, PolyASE testes whether the dominant isoform was consistent across haplotypes. Within each syntenic group, we identified the most expressed haplotype and designated its top two isoforms as “major” and “minor” references. Isoforms were matched across haplotypes using structural similarity scores based on exon count and lengths (60% weight) and intron lengths (40% weight), with differences under 10 bp considered identical. PolyASE tests the major isoform if present across all haplotypes, otherwise the minor isoform. Pairwise comparisons between the reference and each non-reference haplotype were performed.

All likelihood ratio tests used the betabinom_lr_test() function from isotools v2.0.0 (Bi et al., 2025; Lienhard et al., 2023). P-values were corrected for multiple testing using statsmodels.stats.multitest.multipletests() with method = “fdr_bh” (Benjamini & Hochberg, 1995). By default, genes where at least one allele had FDR < 0.05 and a mean ratio difference > 0.1 from the expected ratio were considered to have allelic imbalance or significant difference in allelic expression between conditions. For isoform-level analysis, genes where at least one isoform had FDR < 0.05 and mean isoform ratio differences exceeding 0.2 were classified as having differential isoform usage or haplotype-specific differential isoforms.

For visualization, counts per million (CPM) are calculated from total unique counts per sample and plotted using RNApysoforms v1.2.12 (Aguzzoli Heberle et al., 2024) to compare isoform usage and allele-specific structures.

### Application to Crops Datasets

Sources of all genome annotations and assemblies and long-read RNA-seq datasets used can be found in Supplemental Table S1. Organellar genomes were appended to the respective assemblies, and prefixes were added to gene identifiers to distinguish haplotypes.

Annotation transfer with Liftoff was run independently for each chromosome and haplotype with parameters “-copies -a 0.9 -s 0.9” for rice ‘Nipponbare’ and potato ‘Atlantic’. The Syntelogfinder pipeline v1.0.0 was run on the liftoff annotation of rice ‘Nipponbare’ and potato ‘Atlantic’ and the original annotations of potato ‘Atlantic’, sweet potato ‘Beauregard’ (Wu et al., 2025) and wheat ‘Aikang 58’ (Jia et al., 2023), with reference genome and ploidy of the respective organism.

For *Oryza sativa ssp. japonica ‘*Nipponbare’, a publicly available ONT long-read RNA-seq dataset of rice tillers at four different developmental stages (L, E, M, S) (University, 2025) with three biological replicates was obtained. We compared leaf (L) and early tillering (E) stages. For autotetraploid potato ‘Atlantic’ (Hoopes et al., 2022), a publicly available ONT long-read RNA-seq dataset with five biological replicates from leaf and tuber tissues was obtained.

Both rice and potato long-read RNA-seq samples were processed with the longrnaseq pipeline v1.0.0 using Nextflow v25.04.3 and flag “--technology ONT”. Downsampling rates performed to keep BAM file sizes manageable for visualization were 0.99 for potato and 0.1 for rice.

For gene-level allelic expression analysis with PolyASE in rice and potato, the following settings were used. Only syntenic groups where all replicates had >20 unique reads were retained and single-copy syntelogs present on all haplotypes were selected. For testing allelic differences within conditions, genes were further filtered to retain only those with the same number of isoforms across haplotypes, <0.25 multimapping ratio, and equal CDS lengths.

For isoform-level analysis, only alleles with a minimum expression of 5 uniquely mapping reads in all replicates were retained. Genes with gene-level transcript ambiguity exceeding 0.25 were excluded from the analysis. For testing differences in major isoforms between haplotypes, a minimum similarity of 0.9 was required to match transcripts between haplotypes. For statistical tests default cutoffs (FDR < 0.05 and min ratio difference > 0.1) were used.

## Use Cases

We tested the Syntelogfinder approach on four crops of different ploidies: diploid rice, tetraploid potato, hexaploid sweet potato, and wheat. For rice and potato, for which ONT long-read RNA-seq datasets were available, we tested the end-to-end framework for analyzing allele-specific gene and isoform expression. For rice, we used samples from leaf and early tillering developmental stages, and for potato, we used samples from tubers and leaf tissue.

### Identification of Syntenic Genes

We tested the framework for identification of syntenic relationships on phased assemblies on diploid rice ‘Nipponbare’ (Shang et al., 2023), autotetraploid potato ‘Atlantic’ (Hoopes et al., 2022), allohexaploid sweet potato ‘Beauregard’ (Wu et al., 2025) and allohexaploid wheat ‘Aikang 58’ (Jia et al., 2023). Across all assemblies, a large portion of genes were identified as single-copy syntelogs on each haplotype: 85% in diploid rice (Fig. 2 A), 26% in tetraploid potato (Fig. 2 B), 47% in hexaploid sweet potato, and 44% in hexaploid wheat (Supplemental Fig. S1 A, B).

**Figure 2:**
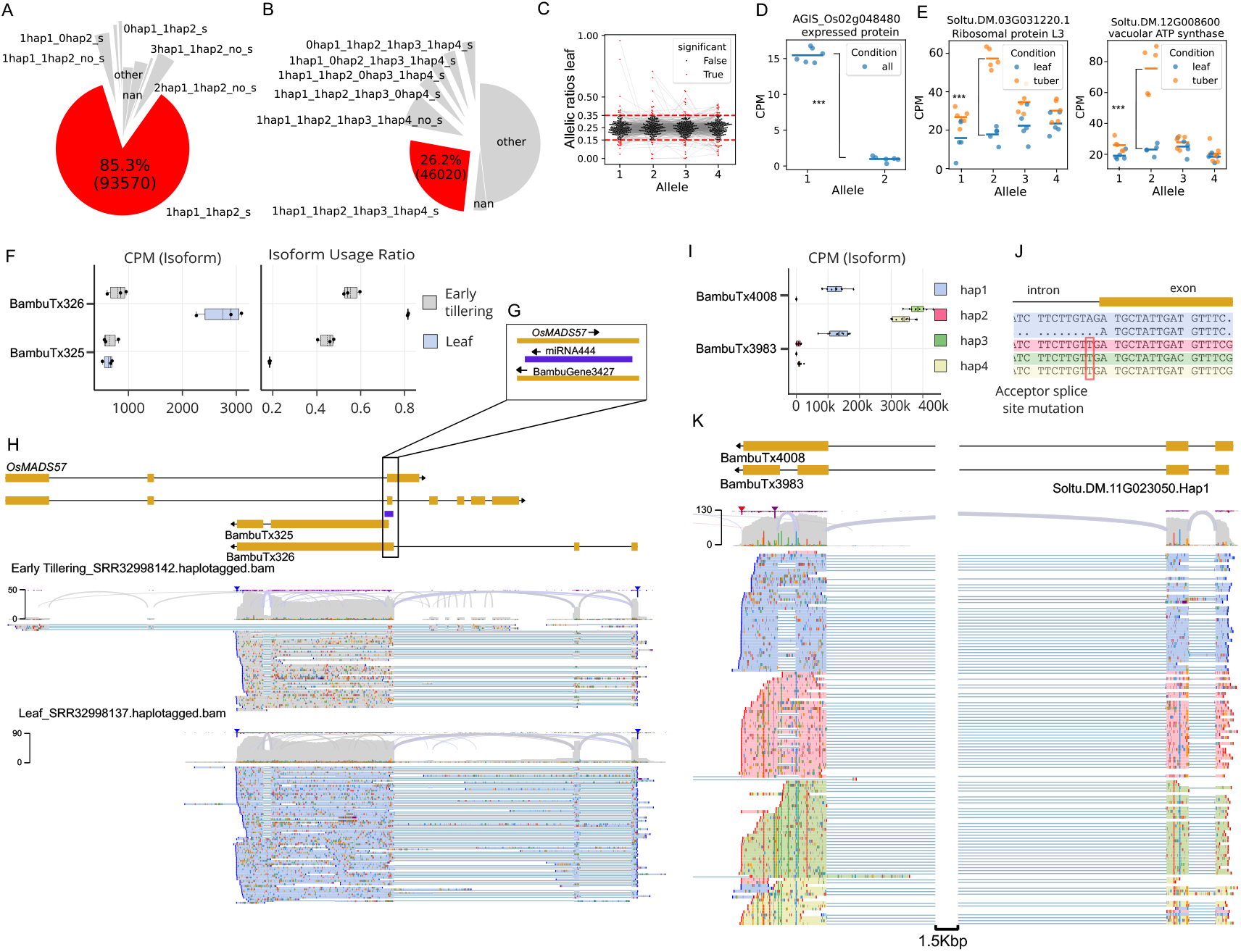
Analysis results of end-to-end framework for allele-specific and isoform expression analysis. Pie charts showing the proportion of genes assigned to syntenic groups for A) rice ‘Nipponbare’ and B) potato ‘Atlantic’. Red indicates single-copy syntelogs, which are present on each haplotype with no gene duplications within any haplotype. Other categories follow the naming convention: [copy number]_hap[haplotype ID]_[synteny status], where ‘_s’ indicates syntenic gene order and ‘_no_s’ indicates non-syntenic genes. C) Swarm plot of allelic ratios for syntelogs in potato leaf samples. Alleles with FDR < 0.05 and mean allelic ratio > 0.35 or < 0.15 are shown in red. Alleles per syntelog are connected with light-gray lines D) Scatter plot of counts per million (CPMs) for rice genes with allelic imbalance. E) Scatter plots of CPMs for potato genes with differences in allelic expression between tuber and leaf tissue. Horizontal lines indicate the mean of biological replicates. F) Box plot of isoform CPMs and isoform usage ratios for the newly annotated rice gene BambuGene3427 with putative differential isoform expression. G) Overlap of the newly identified gene BambuGene3427 with miRNA444 and *OsMADS57*. H) Screenshot of JBrowse2 showing alignment of long-read RNA-seq from L and E samples and structures of newly identified isoforms. I) Box plot of CPMs for isoforms with different major isoform expression across haplotypes in potato. J) Multiple sequence alignment of haplotypes at the splice site with acceptor splice site mutation in haplotypes 2, 3, and 4. K) Screenshot of JBrowse2 showing alignment of long-read RNA-seq sorted and colored by haplotype.

For potato ‘Atlantic’, the original gene annotations of syntelogs exhibit large length differences due to inconsistencies in UTR annotations (Supplemental Fig. S2 C), that introduces a bias in transcript quantification (Nolte et al., 2025). To resolve this bias, either the coding sequences (CDS) can be used as a mapping reference, or gene annotations can be unified between haplotypes using Liftoff. The majority of syntelogs had more than 20% length differences between the longest and the shortest gene model on different haplotypes (Supplemental Fig. S2 C). We mitigated the bias by lifting over the reference double-monoploid’s Unitato gene models (Zagorščak et al., 2024) to the individual haplotypes of the phased assembly using Liftoff. Thereafter, most of the syntelogs showed no length difference between haplotypes (Supplemental Fig. S2 D). We observed similar annotation length bias in the original annotations of wheat and sweet potato (Supplemental Fig. S2 A, B). In the diploid rice gene annotation, which was obtained by lifting over gene annotations from a haploid ‘Nipponbare’ assembly (Shang et al., 2023), syntelogs showed no length difference between haplotypes (Supplemental Fig. S2 E).

### Long-Read RNA-seq QC and Identification of Novel Transcripts

Further analysis was performed in rice and potato for which long-read RNA-seq datasets were available. Quality metrics of raw reads were consistent across samples (Supplementary Table S2). Error rates averaged 5.5% and 7.2% and mapping rates averaged 98% and 82% for rice and potato, respectively. Principal component analysis showed clear separation of samples by tissue in potato, and developmental stage in rice (Supplemental Fig. S3 A, B).

Bambu identified 1521 and 3363 novel genes; and 3278 and 7878 novel transcripts in the rice and potato datasets, respectively. By lifting over these novel transcripts to other haplotypes, 1326 (572) additional transcripts (genes) were added to rice and 8627 (3446) to potato annotation. SQANTI-reads analysis confirmed that reads of all samples exhibited comparable proportions of transcript structures classifications. Most reads were classified as ‘full-splice-matches’ (rice 56-70%, potato 50-70%, Supplemental Fig. S4 A, B). Most of the remaining reads were assigned as ‘incomplete-splice-matches’ and ‘novel-in-catalog’.

### Allelic Expression Quantification

Transcript-level counts were aggregated at the gene-level to assess allelic imbalance between haplotypes and conditions, while transcript counts were used to evaluate differential isoform usage and structural variation between alleles.

#### Gene-Level *Cis*-Regulatory Differences

To investigate *cis*-regulatory variation, we compared expression levels of syntelogs across haplotypes within the same condition or tissue.

In rice ‘Nipponbare’, we tested for imbalance in allelic expression in the leaf developmental stage. The highly inbred rice cultivar exhibited minimal allelic variation, with only 97 of 46,599 syntelogs containing allelic variants (Supplemental Fig. S5). After filtering, only two genes (AGIS_Os12g030940 and AGIS_Os02g048480) were tested, both showing haplotype 1-dominant expression (Fig. 2 D). For AGIS_Os02g048480, read alignment inspection revealed expression exclusively from haplotype 1 (Supplemental Fig. S6), suggesting transcriptional silencing of haplotype 2, which contains two 3’ UTR SNPs and a one-base-pair deletion. No promoter variation was detected within 2kb upstream of the start codon, suggesting epigenetic regulation (e.g. DNA methylation) or distal regulatory control.

In ‘Atlantic’ potato, which is highly heterozygous, we tested for imbalanced expression between haplotypes within the leaf samples. After filtering for length equality, SNP presence and expression level, 211 out of 11,505 syntenic gene groups were tested, with 59 showing significant allelic imbalance at FDR < 0.05 and minimum mean allelic ratio difference > 0.1 (Supplemental Table S3). All haplotypes showed similar proportions of balanced and unbalanced alleles (Fig. 2 C). These results demonstrate how our long-read RNA-seq framework facilitates identification of alleles with imbalanced expression.

#### Gene-Level *Trans*-Regulatory Differences

Since *cis*-regulatory elements associated with each haplotype remain fixed, condition-dependent changes in allelic expression can be caused by *trans*-regulatory factors that vary in abundance across experimental conditions. In rice ‘Nipponbare’, none of the tested syntelogs showed differential allelic expression between the leaf and early tillering development stage.

To detect *trans*-regulatory variation in potato ‘Atlantic’, we tested for differential allelic expression between leaf and tuber tissues. Of 342 syntelogs tested, 165 showed significant differences in the relative expression of at least one allele between tissues at FDR < 0.05, and minimum mean ratio difference > 0.1 between conditions (Supplemental Table S4). For instance, two genes showed tissue-specific imbalance, with haplotype 2 exhibiting higher tuber expression for both a ribosomal protein L3 (ATL_Soltu.DM.12G008600) and vacuolar ATP synthase gene (ATL_Soltu.DM.03G031220) (Fig. 2 E), suggesting that promoters of these genes in haplotype 2 contain a putative tuber-specific transcription factor binding site.

#### Differential Isoform Usage Between Conditions

We tested whether the relative expression of gene isoforms changes between conditions, which can indicate differences in isoform function or condition-specific splicing regulation.

In rice, we identified differential isoform usage of a newly annotated gene (BambuGene3427) with two isoforms (BambuTx326 and BambuTx325). Isoform BambuTx325 showed consistent expression levels between developmental stages, whereas BambuTx326, with a retained intron, is more expressed in the leaf stage (Fig. 2 F). This Bambu-identified gene is predicted to be a non-coding RNA, partially overlapping the tillering regulation gene *OsMADS57* (Fig. 2 H) on the opposite DNA strand. The overlapping region includes a sequence with 142 bp overlap to rice pre-miR444 (Fig. 2 G) suggesting this gene encodes miR444 (Supplemental Fig. S7), which was shown to suppress *OsMADS57* and control tillering (Guo et al., 2013). Notably, *OsMADS57* and the pre-miRNA are present only on haplotype 1. This example demonstrates that our pipeline can identify novel genes and transcripts with differential isoform usage that potentially have functional implications.

#### Differences in Isoform Structure Between Haplotypes

Long-read RNA-seq allows haplotype-resolved measurement of allelic isoform expression and determining whether the predominant isoform expressed from each haplotype shares the same exon-intron structure. The PolyASE package identifies matching isoforms between haplotypes by comparing exon-intron architectures. In potato, we identified 3 out of 228 groups of single-copy syntelogs present once on each haplotype, where the major expressed isoform differed structurally between haplotypes.

One such gene is Soltu.DM.11G023050, for which we identified three isoforms (two of them novel ones) showing differential isoform usage (Fig. 2 I). On haplotype 1, BambuTx4008 and BambuTx3983 were equally expressed, whereas on haplotypes 2, 3, and 4, almost exclusively BambuTx4008 was expressed. This isoform contains a retained intron compared to BambuTx3983 (Fig. 2 K). Multiple sequence alignment revealed that haplotypes 2, 3, and 4 share a mutation in the splice acceptor site (AG → TG), likely causing the intron retention (Fig. 2 J). This example shows that our framework can be effectively used to identify haplotype-specific variations that alter splicing patterns.

## Discussion

We present a computational pipeline specifically designed for allele-specific gene and isoform expression analysis in polyploid organisms using long-read RNA-seq data. Our framework works with datasets of any ploidy level, thus it can be applied to any species with a phased reference genome.

Although pipelines similar to parts of our framework exist, they do not perform haplotype-level quantification of transcripts and cannot handle polyploid genomes. nf-nanoseq (Harshil Patel et al., 2023) performs quantification of known and novel isoforms from ONT long-read RNA-seq, though unlike our framework, it does not process PacBio long-read RNA-seq or include SQANTI3-based quality assessment. TappAS (De La Fuente et al., 2020) conducts differential isoform usage analysis of novel isoforms using short-or long-read RNA-seq, but is not suitable for phased polyploid genomes.

Several challenges remain for the analysis of allelic expression in polyploids. Independent novel transcript discovery on each haplotype creates annotation inconsistencies that complicate comparative isoform analysis. Our implementation using Liftoff partially addresses this issue, although complete resolution would require improved annotation standardization across haplotypes. Furthermore, interpreting the causes of allelic expression differences is challenging, as the mechanisms of transcriptional regulation are still poorly understood (Zeitlinger et al., 2024), thus requiring confirmatory wet lab experiments to identify causal regulatory differences.

Pipeline validation and benchmarking was challenging because publicly available FAIR-compliant long-read RNA-seq datasets with several biological replicates, paired with well-annotated phased genomes, remain scarce. However, we expect that with higher adoption, long reads will replace short-read RNA-seq for haplotype-level expression analysis in polyploids. Although optimal depth will depend on ploidy level and allelic complexity, we recommend an average sequencing depth of ∼100× per allele to ensure reliable allele-specific expression estimates in polyploids.

In future releases, we plan to expand the framework to handle polyploid pangenomes with the aim of supporting comparison of allele expression across cultivars and identifying new breeding targets through pantranscriptomic analysis. Enhanced statistical methods could address genes with high ambiguity and copy number variation. In addition, ongoing advancements in sequencing technologies and analytical tools are likely to improve novel isoform discovery and quantification using long-read RNA-seq.

## Data Availability

Liftoff gene annotations, including novel genes and isoforms, for potato ‘Atlantic’ and rice ‘Nipponbare’ used in this study, are available on Zenodo. In addition, test data is provided on Zenodo for each component of the framework and data to run the complete PolyASE workflow for potato ‘Atlantic’ samples. The longrnaseq pipeline is available at https://github.com/NIB-SI/longrnaseq (doi: https://doi.org/10.5281/zenodo.17631222). The Syntelogfinder pipeline is available at https://github.com/NIB-SI/syntelogfinder (doi: https://doi.org/10.5281/zenodo.17627950). The PolyASE Python package is available at https://pypi.org/project/polyase/ (doi: https://doi.org/10.5281/zenodo.17631236). Documentation and tutorials are available at https://polyase.readthedocs.io/en/latest/index.html.

The long-read RNA-seq is publicly available at NCBI SRA with the following identifiers: *S. tuberosum* ‘Atlantic’ leaf and tuber (SRR14993892 - SRR14993895, SRR14996168, SRR14995031-SRR14995034, SRR14995933) and *O. sativa* ‘Nipponbare’ tiller axillary buds (SRR32998137 and SRR32998141-SRR32998146).

### Runtime Metrics

Minimum recommended hardware: 24 CPU cores, 100 GB RAM, 400 GB storage, and Singularity (Kurtzer et al., 2017), Docker or Conda.

- Syntelogfinder: (30 cores): ∼1 hour runtime
- Long-read RNA-seq pipeline (60 CPU cores): ∼4 hours runtime, 210 GB peak memory (centrifuge), 290-360 GB storage
- Potato dataset: 10 samples, 41.1M reads, 2.1 Gb genome
- Rice dataset: 8 samples, 41.7M reads, 702 Mb genome
- PolyASE analysis: ∼5 minutes

## Supplementary Figures

Figure S1: Pie charts of syntenic groups of genes

Figure S2: Length differences between syntelogs

Figure S3: PCA plots of transcript counts per sample

Figure S4: SQANTI-reads categories per sample

Figure S5: Barplot of number of syntelogs per identity category for rice

Figure S6: Read alignments to AGIS_Os02g048480 haplotype 2 in rice

Figure S7: Overlap of BambuTx325 and BambuTx326 with pre-miRNA *Os*mir444d

## Supplementary Tables

Table S1: Availability of reference files and sample information of long-read RNA-seq samples

Table S2: Mapping statistics of long-read RNA-seq of rice ‘Nipponbare’ and potato ‘Atlantic’

Table S3: Results of allele-specific expression analysis within control conditions for potato ‘Atlantic’

Table S4: Results of allele-specific expression analysis between tuber and leaf tissue for potato ‘Atlantic’

## Author Contributions Statement

**Nadja Nolte:** Conceptualization (Equal), Formal analysis (Lead), Investigation (Lead), Methodology (Lead), Software (Lead), Visualization (Lead), Writing - original draft (Lead), Writing - review & editing (Equal). **Kristina Gruden:** Funding acquisition (Equal), Writing - review & editing (Equal). **Marko Petek:** Conceptualization (Equal), Funding acquisition (Equal), Supervision (Lead), Writing - review & editing (Equal).

## Acknowledgements

This work was supported by the European Union’s Horizon 2020 research and innovation programme under the Marie Skłodowska-Curie Actions Doctoral Network “LongTREC” [grant number 101072892]; and the Slovenian Research and Innovation Agency grant agreements [grant numbers P4-0463, P4–0431].

The authors gratefully acknowledge Academic and Research Network of Slovenia (ARNES) for providing computing resources of the Arnes HPC cluster. We also thank all the LongTREC network for discussion on long-read RNA-seq mapping approaches and tools for isoform identification.

