## Supplemental Figures S1 - S7 for "LongPolyASE: An end-to-end framework for allele-specific gene and isoform analysis in polyploids using long-read RNA-seq"

### Supplementary Figures

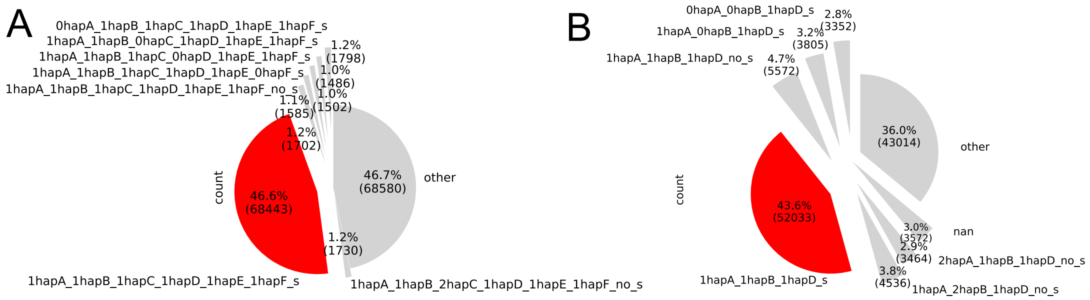


**Figure S1**: Pie chart showing the number of genes per syntenic group for A) hexaploid sweet potato ‘Beauregard’ and B) ABC subgenomes of hexaploid wheat ‘Aikang 58’. In red: syntenic category where syntelogs are once on each haplotype. ‘_s’: syntenic gene order, ‘_no_s’: non syntenic, number before haplotype (hap) indicates copy number.

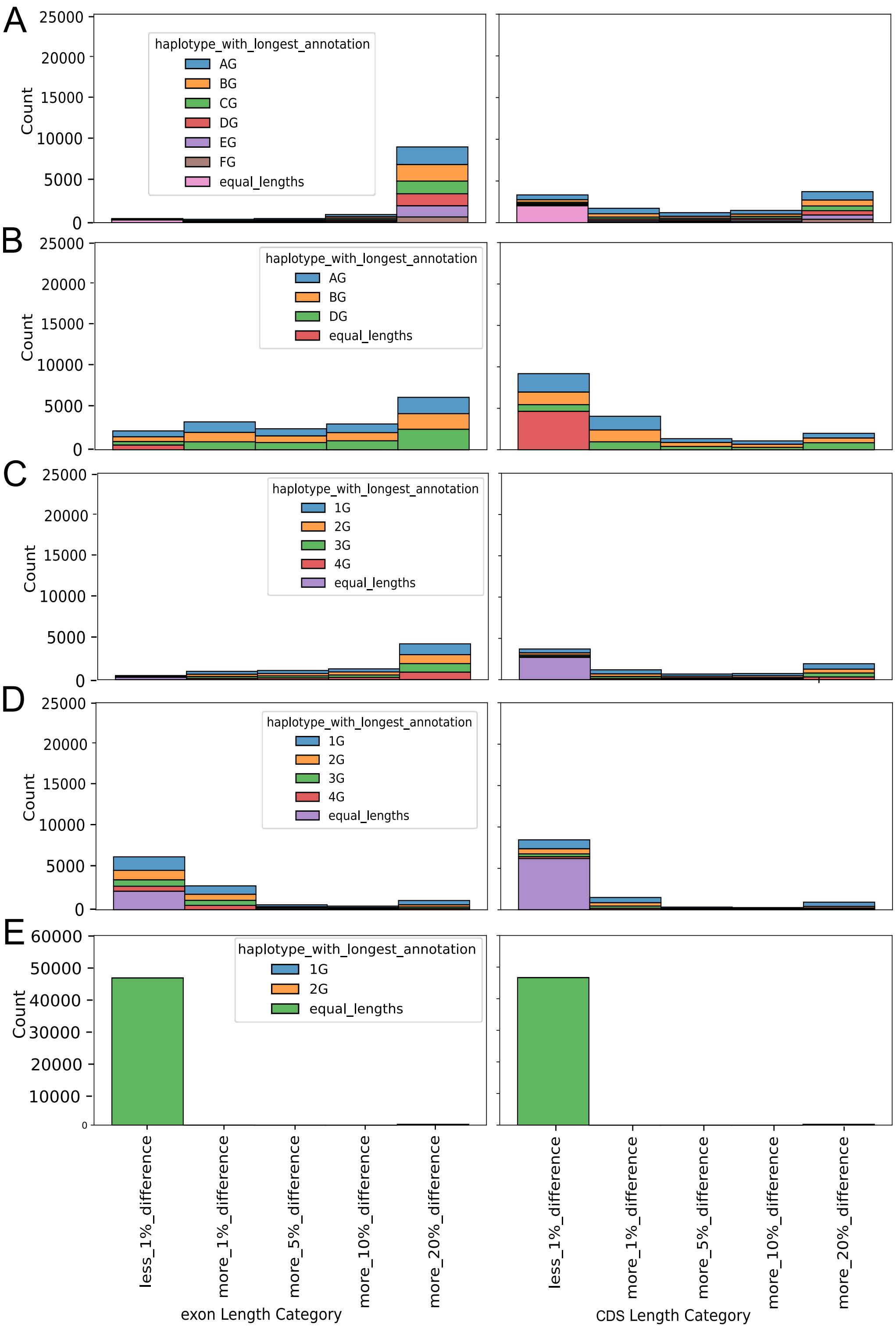


**Figure S2**: Barplot showing number of transcripts in each length category. Longest transcripts per gene in each syntenic group were categorized based on the differences between longest and shortest transcript for A) hexaploid assebmly of sweet potato 'Beauregard' B) triploid subgenome assembly of wheat 'Aikang 58' C) tetraploid assembly of potato 'Atlantic' with original gene annotation D) tetraploid assembly potato 'Atlantic' with Liftoff gene annotation from double monoploid potato and (E) diploid assembly of rice 'Nipponbare' with Liftoff gene annotation.

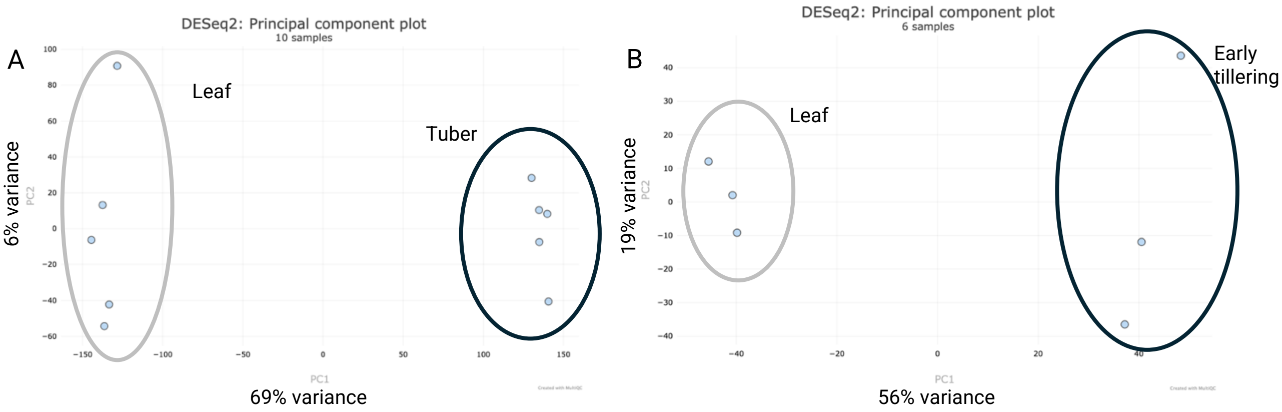


**Figure S3**: PCA plots of ONT long-read RNA-seq samples for (A) rice 'Nipponbare' and (B) potato 'Atlantic'. Normalized transcript counts were used to generate the plots. Circles enclose biological replicates for leaf and tuber samples in potato, and leaf and early tillering developmental stages in rice.

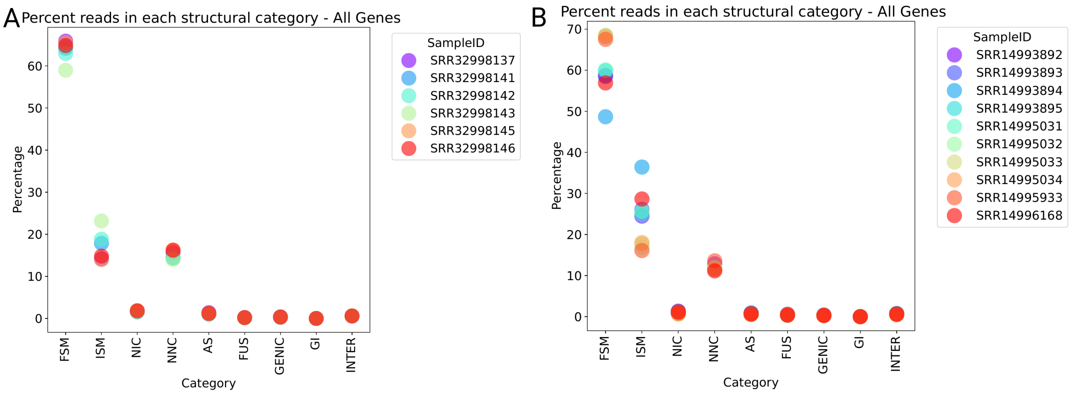


**Figure S4**: Sqanti-reads analysis output with assignment of reads to structural categories for (A) rice 'Nipponbare' and (B) potato 'Atlantic'. FSM: full-splice-match, ISM: incomplete-splice-match, NIC: novel-in-catalog, NNC: novel-not-in-catalog, AS: antisense, FUS: Fustion gene, GENIC: genic genomic, GI: genic intron, INTER: intergenic.

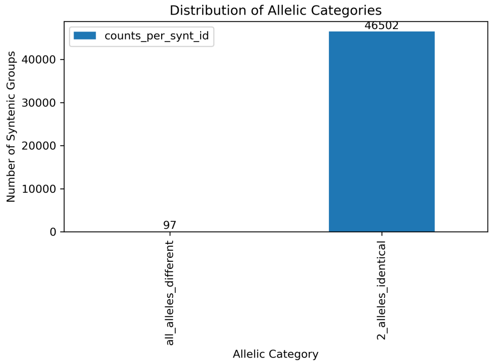


**Figure S5**: Barplot of number of genes per Identity category for one-to-one syntelogs in rice 'Nipponbare'.

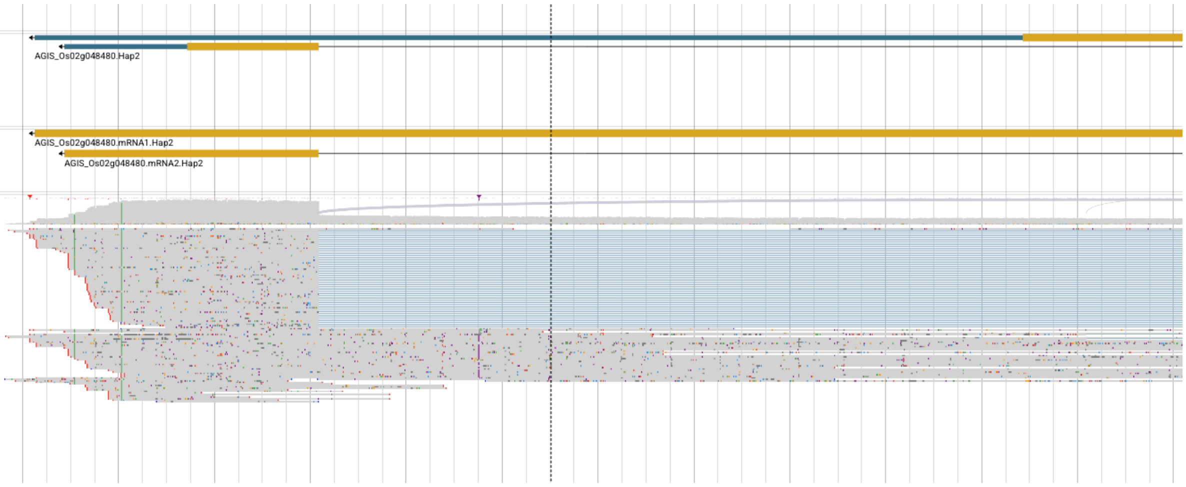


**Figure S6**: Screenshot of long-read RNA-seq reads aligning to AGIS_Os02g048480.Hap2 in rice 'Nipponbare'.

Oryza sativa miR444d stem-loop (osa-MIR444d)

143 nucleotides

```
Query   71  AGUUAUUGCACAUGGUGGCACCAAGCAUGAGGCAACAACUGCAUUACUUGCAAGAAAGGCACAAAAUCAUUAGAUGAUUACUUGUGGCUUU  161
            |||||||||||||||||||||||||||||||||||||||||||||||||||||||||||||||||||||||||||||||||||||||||||
Sbjct    1  AGUUAUUGCACAUGGUGGCACCAAGCAUGAGGCAACAACUGCAUUACUUGCAAGAAAGGCACAAAAUCAUUAGAUGAUUACUUGUGGCUUU  91

Query  162  CUUGCAAGUUGUGCAGUUGCUGCCUCAAGCUUGCUGCCUCCCUCUGCCAAA  212
            |||||||||||||||||||||||||||||||||||||||||||||||||||
Sbjct   92  CUUGCAAGUUGUGCAGUUGCUGCCUCAAGCUUGCUGCCUCCCUCUGCCAAA  142
```

**Figure S7**: Overlap of newly identified transcripts BambuTx325 and BambuTx326 and pre-miRNA of miR444d in rice 'Nipponbare' in miRbase. miRNA44 is highlighted in red.
